# Penetrance of polygenic obesity susceptibility loci across the body mass index distribution: an update on scaling effects

**DOI:** 10.1101/225128

**Authors:** Arkan Abadi, Akram Alyass, Sebastien Robiou du Pont, Ben Bolker, Pardeep Singh, Viswanathan Mohan, Rafael Diaz, James C. Engert, Hertzel C. Gerstein, Sonia S. Anand, David Meyre

## Abstract

A growing number of single nucleotide polymorphisms (SNPs) have been associated with body mass index (BMI) and obesity, but whether the effect of these obesity susceptibility loci is uniform across the BMI distribution remains unclear. We studied the effects of 37 BMI/obesity-associated SNPs in 75,230 adults of European ancestry along BMI percentiles using conditional quantile regression (CQR) and meta-regression (MR) models. The effects of 9 SNPs (24%) increased significantly across the sample BMI distribution including, *FTO* (rs1421085, p=8.69×10^−15^), *PCSK1* (rs6235, p=7.11×10^−06^), *TCF7L2* (rs7903146, p=9.60×10^−06^), *MC4R* (rs11873305, p=5.08×10^−05^), *FANCL* (rs12617233, p=5.30×10^−05^), *GIPR* (rs11672660, p=1.64×^−04^), *MAP2K5* (rs997295, p=3.25×10^−04^), *FTO* (rs6499653, p=6.23×10^−04^) and *NT5C2* (rs3824755, p=7.90×10^−04^). We showed that such increases stem from unadjusted gene interactions that enhanced the effects of SNPs in persons with high BMI. When 125 height-associated were analyzed for comparison, only one (<1%), *IGF1* (rs6219, p=1.80×10^−04^), showed effects that varied significantly across height percentiles. Cumulative gene scores of these SNPs (GS-BMI and GS-Height, respectively) showed that only GS-BMI had effects that increased significantly across the sample distribution (BMI: p=7.03×10^−37^, Height: p=0.499). Overall, these findings underscore the importance of gene-gene and gene-environment interactions in shaping the genetic architecture of BMI and advance a method to detect such interactions using only the sample outcome distribution.

## INTRODUCTION

Obesity is a prominent risk factor for psychological disorders, osteoarthritis, hypertension, type 2 diabetes (T2D), cardiovascular disease, and certain cancers.^1,2^ The rise of obesity coincided with ‘obesogenic’ societal and environmental changes that include excessive consumption of high calorie foods, sedentary lifestyle and urbanization.^2–4^ Genetic factors are also known to play an important role in obesity as 50–80% of body mass index (BMI) variation can be ascribed to genetics (heritability).^5,6^ Moreover, genome wide association studies (GWAS) have identified ~140 polygenic loci that are directly associated with BMI or obesity.^7^

The role of individual and compound gene-environment (GXE) and gene-gene (GXG) interactions in determining BMI has not been fully elucidated. The study of BMI-associated GXG interactions has been impeded by statistical and computational limitations, although promising new approaches have recently been proposed.^8–10^ On the other hand, several lines of evidence suggest that GXE interactions may play an important role in shaping BMI. First, estimates of the heritability of BMI are influenced by environmental exposures.^11^ One study reported that the heritability of BMI is increased in persons born after the obesogenic transition, while another reported that the heritability of BMI is correlated with the population prevalence of obesity.^12,13^ More recently, the cumulative gene score from 29 BMI-associated SNPs showed a positive interaction effect with birth year.^14^ Interactions between the genetic determinants of BMI and obesogenic environmental factors readily explains why both estimates of BMI heritability and cumulative SNP effects are enhanced in permissive environments. Second, specific interactions between BMI-associated SNPs and environmental factors have been documented.^11^ Physical activity and energy intake have been reported to modify the effects of SNPs within the fat mass and obesity-associated (*FTO*) gene.^15–19^ Importantly, *FTO* (rs1421085) has been shown to jointly interact with diet, physical activity, salt and alcohol consumption and sleep duration.^20^ Thus, a subset of genetic variants may affect BMI through a mixture of direct effects and compound interactions. As such, investigating individual environmental factors may not capture the full range of environmental modification for a given SNP^21,22^

In this report, we advanced a statistical framework to assess the effects of single and mixed GXE and GXG interactions on the association between SNPs and BMI. Specifically, conditional quantile regression (CQR) was applied to investigate the effects of 37 BMI/obesity-associated SNPs at multiple percentiles of the sample BMI distribution in 75,230 adults of European ancestry (EA).^23,24^ Variability in SNP effects across these BMI percentiles was demonstrated to result from unadjusted interactions and was modeled using meta-regression (MR).^25,26^ In this way, CQR and MR were used to collect evidence of unadjusted interactions directly from the sample distribution of BMI absent measures of specific environmental factors. A secondary analysis on 125 established height-associated SNPs is also included for comparison.

## SUBJECTS AND METHODS

### Participants and Phenotypes

The sample population included participants from the following studies: ARIC (phs000280.v3.p1), CARDIA (phs000285.v3.p2), CHS (phs000287.v6.p1), EpiDREAM, the Framingham cohort (phs000007.v29.p10), MESA (phs000209.v13.p3), COPD (phs000179.v5.p2), eMERGE II (phs000888.v1.p1) and the WHI (phs000200.v10.p3). Measurements collected from participants below age 18 or above the age of 92 were excluded (<1%). For studies with repeated measures across multiple time points or visits, the median height and the median weight was extracted along with the corresponding age at these median values. BMI was calculated by dividing the median weight (kg) by the square of the average measures of height (m). Diabetic status was indicated by one of the following criteria: (1) physician report or self-report of physician-diagnosis, (2) report taking diabetes medication, (3) fasting plasma glucose ≥ 126 mg/dL (7mM), or (4) 2hr glucose ≥ 200 mg/dl (11mM) during an oral glucose tolerance test (OGTT).^27^ Obesity categories including, normal weight (NW), overweight (OW), as well as obesity classes I, II and III (Ob-l, Ob-ll, and Ob-Ill, respectively) were specified according to WHO guidelines.^28^ Analyses were restricted to participants of European ancestry (EA, self-reported) with a combined sample size of N=75,230. Summary statistics are presented Table S1. This project was approved by local ethics committee (Hamilton Integrated Research Ethics Board-HiREB) and participant-level data access was granted through the database of Genotypes and Phenotypes (dbGaP) following approval by study-specific Data Access Committee (DAC). All analyses are consistent with study-specific Data Use Certifications (DUC).

### Sample Quality Control (QC)

Detailed genotyping procedures for EpiDREAM and studies from the Candidate Gene Association Resource (CARe) project including, ARIC (phs000557.v2.p1), CARDIA (phs000613.v1.p2), CHS (phs000377.v4.p1), the Framingham Cohort (phs000282.v17.p10) and MESA (phs000283.v7.p3) are presented elsewhere.^29,30^ Genotyping was performed using the gene centric HumanCVD Genotyping BeadChip with 49,320 markers concentrated in ~ 2,100 loci that are related to metabolism and cardiovascular disease.^31^ This limited scope of analysis was motivated by access to a greater sample size, as well as the high computational cost of fitting CQR models across percentiles the sample outcome distribution. Samples with sex-discordance, array-wide call rate below 95–98%, and/or average heterozygosity beyond 3 standard deviations of the mean heterozygosity, were removed.^32,33^ Family members were defined by identity by descent (IBD) (PI HAT) above 0.5 and those with lower call rate were removed so that only one member of each family group was retained for analysis (Table S2). Samples from the COPDGene (phs000765.v1.p2) study were genotyped using the lllumina HumanHap550 (v3) genotyping BeadChip (lllumina Inc., San Diego, CA, USA) with 561,466 markers and QC procedures were performed as above except that cryptic relatedness was defined by IBD PI HAT > 0.1875.^34,35^ Genotypes from the WHI study (phs000746.v1.p3) and eMERGE II (phs000888.v1.p1) were comprised of an imputed dataset and samples from related/duplicate participants were removed.

Analyses of the WHI dataset were conducted on each sub-study (WHIMS, GARNET, HIPFX, MOPMAP and GECCO). A summary of sample QC, along with a complete list of datasets (accession #) and additional details on these studies is provided in Table S2.

### SNP Selection and Marker QC

SNPs that have previously been associated with BMI, obesity and height were identified by searching the genome-wide association study (GWAS) Catalog, GIANT Consortium data files, and screening the literature.^36–39^ Literature screening was conducted independently by A.A. and D.M. to maximize SNP attainment. For GWAS SNPs, only associations with p<5×10^−08^ were considered. These SNPs were sorted into correlated linkage disequilibrium blocks (LD, R^2^ > 0.1) based on genomic sequences from EA populations (Phase 3, 1000 Genome Project) and the strongest association SNP on the HumanCVD Genotyping BeadChip was selected.^31,40^ Proxy SNPs (LD, R^2^ > 0.9) were identified for SNPs in LD blocks not represented on the array. Thus, 39 BMI/obesity- and 129 height-associated SNPs were identified. For studies using different genotyping platforms, the original association SNPs (39 BMI/obesity and 129 height) were screened and proxied as above on each genotyping platform. SNPs that mapped to the same gene were screened jointly using conditional regression analysis to test for independent associations with quantitative traits (BMI or height) and only SNPs that maintained associations were retained.^41^ However SNPs in *FTO* (rs1421085 and rs6499653) and *PCSK1* (rs6232 and s6235) were exempted from exclusion due to prior evidence in the literature of independent associations with BMI/obesity.^42–44^ In total, 37 BMI/obesity- and 125 height-associated independent SNPs were identified and selected for further analysis. SNP call rate, minor allele frequency (MAF) and exact tests of Hardy-Weinberg equilibrium (HWE) in EA populations are presented in Tables S3-4. Within each study, SNPs with a call rate below 90% or HWE p-value<1×10^−06^ were excluded from analysis. In addition, only SNPs imputed with high quality were retained for analysis (R^2^ > 0.7 for WHI and info score > 0.7 for eMERGE II).^45^ SNP genotypes were encoded per the trait increasing effect alleles and modeled additively for individual analyses.

### Gene Scores

The cumulative gene score (GS) was calculated for all BMI/obesity- and height-associated SNPs (GS-BMI and GS-Height, respectively). An un-weighted GS was utilized because weights can be biased and context dependent.^46,47^ No GS was calculated for participants with more than 10% missing genotypes, otherwise missing SNP genotypes were imputed using the arithmetic average genotype at each missing SNP. In addition to BMI, *GIPR* (rs10423928, LD R^2^=1 with rs11672660 in EA), *TCF7L2* (rs7903146), *TOMM40/APOE* (rs2075650), *HMGCR* (rs4604177, LD R^2^=0.63 with rs6453133 in EA), *PCSK1* (rs6235), *CDKAL1* (rs9356744) and *KCNQ1* (rs2283228) have also been associated with several co-morbidities of obesity including glucose homeostasis, T2D, lipid levels and CRP levels.^48–55^ To mitigate potential biases stemming from these comorbidities at higher BMI percentiles, a GS excluding these 7 SNPs was also calculated, GS-BMI (Stringent). Finally, GSs for both BMI and Height were calculated without imputing missing genotypes, GS-BMI (No Imputation) and GS-Height (No Imputation). Testing GS-BMI (Stringent), GS-BMI (No Imputation) and GS-Height (No Imputation) was performed as sensitivity analysis.

### Statistical Analysis

A statistical framework combining CQR and MR was used to model variation in the effects of SNPs under single and mixture GXE and GXG interactions (see Supplemental Note).^24,26^ Like ordinary least square (OLS), CQR models may assume a linear relationship and provide intercept and slope estimates for a series of pre-specified percentiles.^23,24^ Therefore, CQR can be applied to produce a comprehensive evaluation of the effects of a SNP across the sample distribution of a quantitative trait (e.g. BMI or height). A piecewise linear plot for the series of CQR estimates at different percentiles provides a useful visual summary of their variation along the sample distribution.^23,24^ Figure 1 shows a working example of CQR and MR in comparison with OLS using *FTO* (rs1421085) and ARIC CARe.

**Figure 1:**
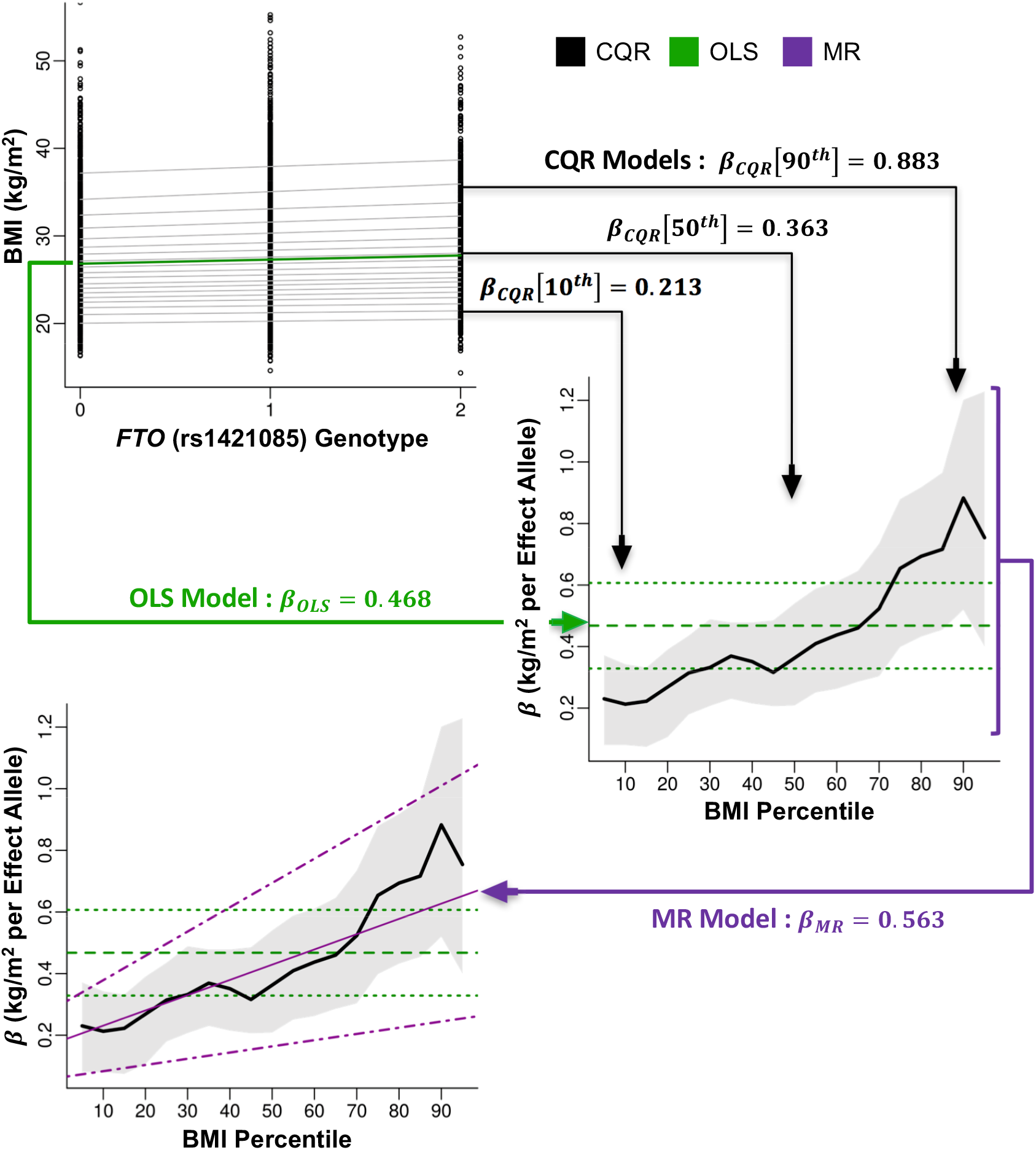
*Working example of conditional quantile regression*. BMI (kg/m^2^) was plotted against the number of effect alleles of *FTO* (rs1421085) in the ARIC CARe study (top-left). An ordinary least squares (OLS) model of the mean effect of this SNP on BMI was plotted (solid-green line). Conditional quantile regression (CQR) models, fitted every 5^th^ percentile of BMI, show the effects of this SNP at these BMI percentiles (solid-grey lines). The slopes (*β_OLS_*, horizontal-dashed-green line; *β_CQR_*, thick-black line; kg/m^2^ per Effect Allele) from these models were then plotted against BMI percentile at which they were fitted (middle-right). 95% confidence intervals for these estimates are also plotted (OLS, horizontal-dotted-green line; CQR, shaded-grey region). The change in CQR estimates across BMI percentiles was modeled using meta-regression (MR). The MR slope (*β_MR_*, kg/m^2^ per Effect Allele per BMI percentile, thin-magenta line) and the 95% confidence intervals (dotdashed-magenta lines) were plotted (bottom-left).

Under conditions where true single and mixed GXE and GXG interactions are unadjusted, SNPs will shift both the location and scale (variance) of the sample outcome distribution (see Supplemental Note).^56^ These shifts in scale result in detectable variations of CQR estimates collected from percentiles across the sample outcome distribution. It follows that CQR estimates for a SNP are constant (i.e. equal) across percentiles if all unadjusted interaction effects are zero. It is important to consider that GXG interactions include non-linear genetic models of effects where the presence of one allele changes the effects of a second allele within the same variant.^57,58^ Thus, the association of SNPs with an outcome under unadjusted interactions essentially reduces to modelling variability in CQR estimates. This can be effectively achieved using MR.^25,26^ In this context, MR is basically a regression model where the CQR estimates from across the sample outcome distribution represent the dependent variable and the percentiles at which these CQR estimates were calculated represent the independent variable (Figure 1). Additional details on CQR and MR as well as an analytic description of this statistical framework and simulations are presented in the Supplemental Note and Figures S1-2.

OLS models were used to verify the associations of SNPs and GSs with BMI/obesity and height in the sample populations included in this study. CQR models were fitted at every 5^th^ percentile of the distribution of BMI and height for each SNP. A total of 10,000 Markov chain marginal bootstrap (MCMB) replicates were used to compute confidence intervals and the cross-percentile variance-covariance matrix for CQR estimates.^59–61^ The proportion of the trait variance explained by GS-BMI and GS-Height in CQR models was also calculated.^62^ Hypothesis test statistics in MR were computed assuming normality to estimate the effects of percentiles on changes in mean CQR estimates for each SNP. The set of percentiles (5^th^ to 95^th^) were re-centered at the 50^th^ percentile so that the intercept of the MR models corresponds to the main effect of the SNP at the median. To compute residuals after adjusting for the median effects of each SNP on BMI, the univariable median effect of each SNP was calculated using CQR at the 50^th^ percentile and the residuals were calculated by subtracting the product of this median effect and the genotype from BMI. SNP effects on variance were estimated using OLS analysis of z^2^.^63^ Lastly, the effects of each SNP and the GS on the risk of specific BMI categories (NW vs. OW, NW vs. Ob-l, NW vs, Ob-ll and NW vs. Ob-Ill) was estimated using logistic regression.

All regression models were performed using one-step individual participant-data meta-analysis (also known as ‘joint-analysis’ or ‘mega-analysis’).^64,65^ This method was justified by access to individual participant data and the fact that CQR analyses refer to the conditional sample distribution.^66^ This means that analyses on separate studies correspond to their conditional distributions and it would not be appropriate to combine them using metaanalysis of their summary statistics. All models were adjusted for age (years), sex (female=0, male=1) and study (factor). Age was modeled quadratically (age and age-squared) for BMI analysis, consistent with previous reports.^14,20^ Analysis of SNPs and the GS with BMI (37 SNPs+GS=38) and height (125 SNPs+GS=126) were subject to multiple testing correction using Bonferroni adjusted p-value thresholds of p<0.05/38=1.32×10^−03^ and p<0.05/126=3.97×10^−04^, respectively.^67^ QC and statistical analyses were conducted using PLINK version 1.90b3.42 and R version 3.3.2.^32,33,68–78^ CQR models were fitted using *quantreg* and MR models were fitted using *metafor.^79,80^* Additional packages used in the analysis include *GenABEL, pracma, doParallel, foreach* and *data.table*.^81–85^

## RESULTS

Figure 1 depicts a step by step analysis of *FTO* (rs1421085) in the ARIC CARe study. In the top left panel, an OLS model (green) is fitted to determine the mean effects of *FTO* genotype on BMI (*β_OLS_*, kg/m^2^ per Effect Allele) and CQR models (grey) are fitted evenly across the sample BMI distribution (every 5^th^ percentile) to determine the effects of *FTO* genotype at each BMI percentile (*β_CQR_*, kg/m^2^ per Effect Allele). In middle right panel, the estimates (*β_OLS_* and *β_CQR_*) and 95% confidence intervals from these models are collected and plotted against the BMI percentile at which they were fitted. In the bottom left panel, MR analysis (magenta) is used to model variation in the CQR estimates across the sample BMI distribution and MR estimates (*β_MR_*, kg/m^2^ per Effect Allele per BMI percentile) along with 95% confidence intervals are plotted. Presenting the results of OLS, CQR and MR in this way is useful for summarizing the purpose of each analysis and contrasting possible differences between them.

Initially, OLS models were fitted for each of 37 BMI/obesity-associated SNPs and all but one was verified to increase BMI in this study sample (Table 1). CQR models were then fitted at regular intervals of the BMI distribution to explore whether the effects of SNPs on BMI varied across the sample distribution (Table S5). CQR estimates for each SNP were plotted against the BMI percentiles at which they were produced to provide a visual summary of the CQR results (Figures 2 and S3). Several SNPs had effects that appeared to increase across the distribution of BMI including, *FTO* (rs1421085), *PCSK1* (rs6235), *TCF7L2* (rs7903146), *MC4R* (rs11873305), *FANCL* (rs12617233), *GIPR* (rs11672660), *MAP2K5* (rs997295), *FTO* (rs6499653) and *NT5C2* (rs3824755).

**Figure 2:**
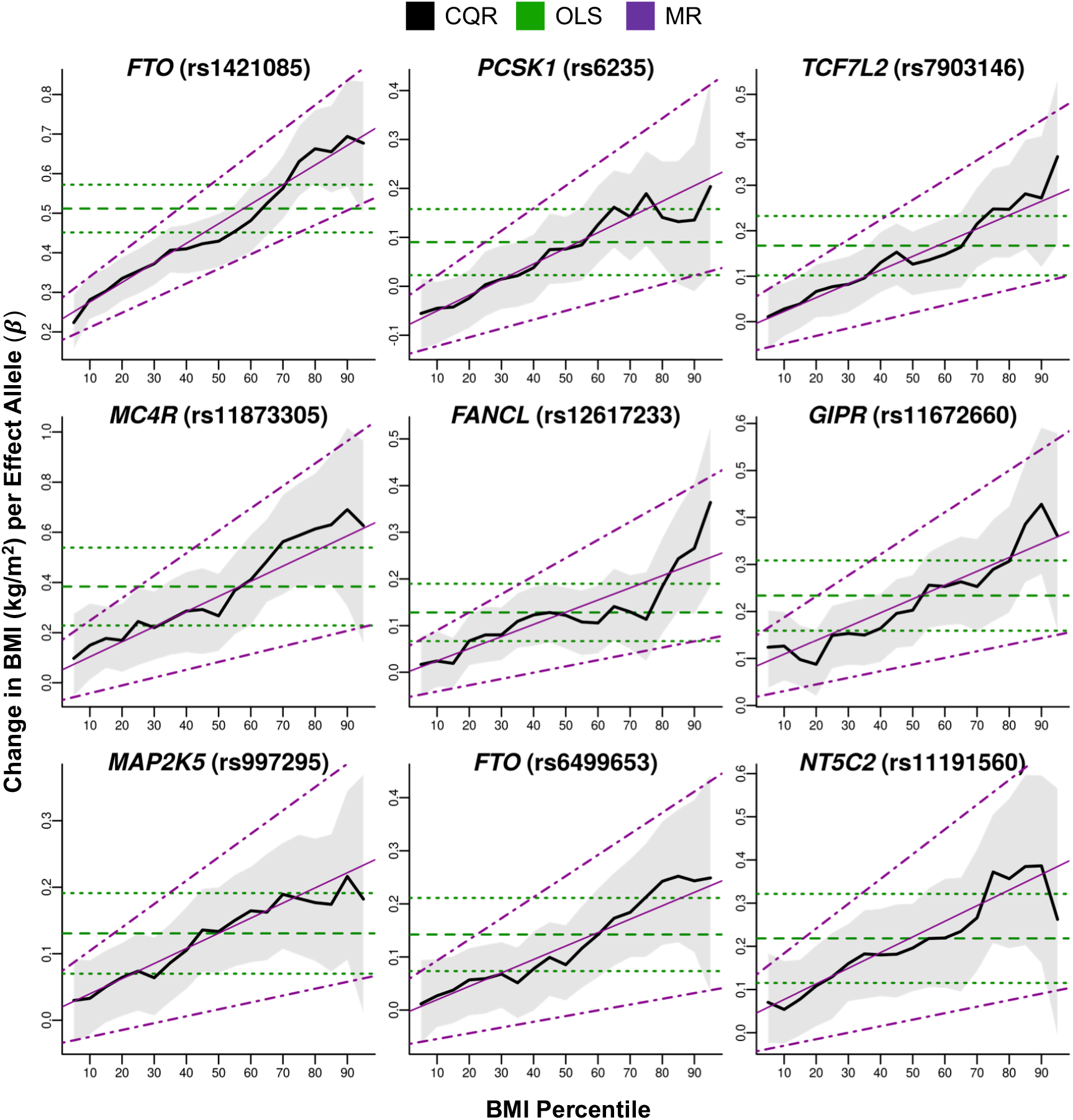
*The effects of BMI/obesity-associated SNPs across the sample BMI distribution*. Conditional quantile regression (CQR) models of BMI/obesity-associated SNPs were fitted every 5^th^ percentile of BMI and adjusted for age, age-squared, sex and study. Estimates of the change in BMI (kg/m^2^) per effect allele (*β_CQR_*) from these models was plotted against the BMI percentile (thick-black line) along with the 95% confidence intervals (shaded-grey region). The results from ordinary least square (OLS) models (*β_OLS_*, kg/m^2^ per Effect Allele, horizontal-dashed-green line) and the 95% confidence intervals (horizontal-dotted-green lines) were also plotted for comparison. The change in CQR estimates across BMI percentiles was modeled using meta-regression (MR) and estimates from MR (*β_MR_*, kg/m^2^ per Effect Allele per BMI percentile, thin-magenta line) and the 95% confidence intervals (dotdashed-magenta lines) were plotted. MR analysis detected significant (p<1.32×10^−03^) increases in the effects of these SNPs across the sample BMI distribution.

**Table 1:**
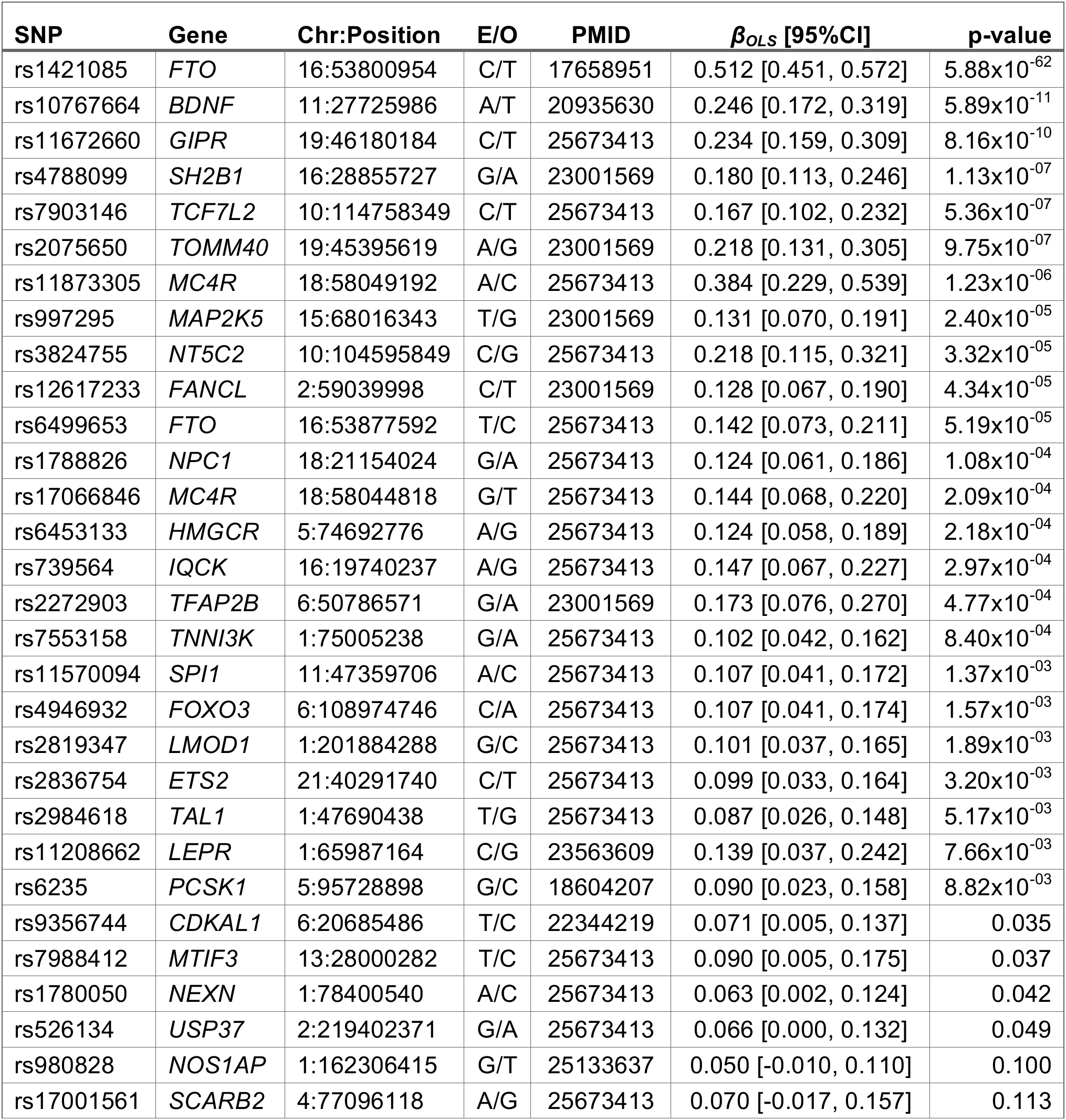
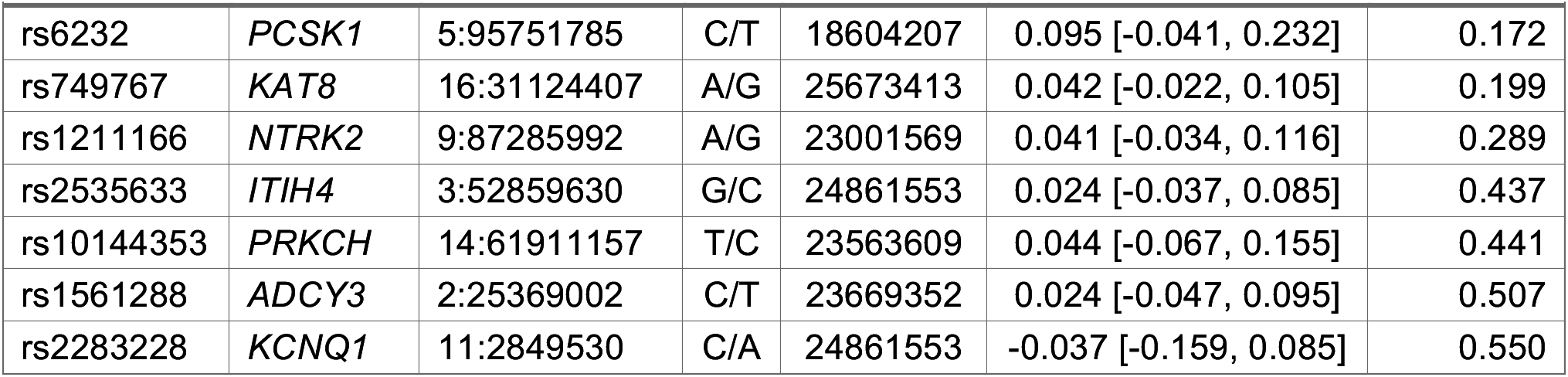
*BMI/Obesity-associated SNP Information and results from ordinary least squares (OLS) models*. 37 BMI/obesity predisposing SNPs were selected for analysis. The Effect / Other (E/0) alleles were based on original discovery studies (PMID), and SNPs were coded by BMI increasing or obesity predisposing alleles. Indicated positions were based on GRCh37 and all alleles were on the positive strand. The association of these SNPs with BMI was assessed using OLS models that were adjusted for age, age-squared, sex and study. *β_OLS_* is the effect size (kg/m^2^ per Effect Allele) and 95%CI are the 95% confidence intervals.

| SNP | Gene | Chr:Position | E/O | PMID | $\beta_{OLS}$ [95%CI] | p-value |
| --- | --- | --- | --- | --- | --- | --- |
| rs1421085 | *FTO* | 16:53800954 | C/T | 17658951 | 0.512 [0.451, 0.572] | 5.88x10<sup>-62</sup> |
| rs10767664 | *BDNF* | 11:27725986 | A/T | 20935630 | 0.246 [0.172, 0.319] | 5.89x10<sup>-11</sup> |
| rs11672660 | *GIPR* | 19:46180184 | C/T | 25673413 | 0.234 [0.159, 0.309] | 8.16x10<sup>-10</sup> |
| rs4788099 | *SH2B1* | 16:28855727 | G/A | 23001569 | 0.180 [0.113, 0.246] | 1.13x10<sup>-07</sup> |
| rs7903146 | *TCF7L2* | 10:114758349 | C/T | 25673413 | 0.167 [0.102, 0.232] | 5.36x10<sup>-07</sup> |
| rs2075650 | *TOMM40* | 19:45395619 | A/G | 23001569 | 0.218 [0.131, 0.305] | 9.75x10<sup>-07</sup> |
| rs11873305 | *MC4R* | 18:58049192 | A/C | 25673413 | 0.384 [0.229, 0.539] | 1.23x10<sup>-06</sup> |
| rs997295 | *MAP2K5* | 15:68016343 | T/G | 23001569 | 0.131 [0.070, 0.191] | 2.40x10<sup>-05</sup> |
| rs3824755 | *NT5C2* | 10:104595849 | C/G | 25673413 | 0.218 [0.115, 0.321] | 3.32x10<sup>-05</sup> |
| rs12617233 | *FANCL* | 2:59039998 | C/T | 23001569 | 0.128 [0.067, 0.190] | 4.34x10<sup>-05</sup> |
| rs6499653 | *FTO* | 16:53877592 | T/C | 25673413 | 0.142 [0.073, 0.211] | 5.19x10<sup>-05</sup> |
| rs1788826 | *NPC1* | 18:21154024 | G/A | 25673413 | 0.124 [0.061, 0.186] | 1.08x10<sup>-04</sup> |
| rs17066846 | *MC4R* | 18:58044818 | G/T | 25673413 | 0.144 [0.068, 0.220] | 2.09x10<sup>-04</sup> |
| rs6453133 | *HMGCR* | 5:74692776 | A/G | 25673413 | 0.124 [0.058, 0.189] | 2.18x10<sup>-04</sup> |
| rs739564 | *IQCK* | 16:19740237 | A/G | 25673413 | 0.147 [0.067, 0.227] | 2.97x10<sup>-04</sup> |
| rs2272903 | *TFAP2B* | 6:50786571 | G/A | 23001569 | 0.173 [0.076, 0.270] | 4.77x10<sup>-04</sup> |
| rs7553158 | *TNNI3K* | 1:75005238 | G/A | 25673413 | 0.102 [0.042, 0.162] | 8.40x10<sup>-04</sup> |
| rs11570094 | *SPI1* | 11:47359706 | A/C | 25673413 | 0.107 [0.041, 0.172] | 1.37x10<sup>-03</sup> |
| rs4946932 | *FOXO3* | 6:108974746 | C/A | 25673413 | 0.107 [0.041, 0.174] | 1.57x10<sup>-03</sup> |
| rs2819347 | *LMOD1* | 1:201884288 | G/C | 25673413 | 0.101 [0.037, 0.165] | 1.89x10<sup>-03</sup> |
| rs2836754 | *ETS2* | 21:40291740 | C/T | 25673413 | 0.099 [0.033, 0.164] | 3.20x10<sup>-03</sup> |
| rs2984618 | *TAL1* | 1:47690438 | T/G | 25673413 | 0.087 [0.026, 0.148] | 5.17x10<sup>-03</sup> |
| rs11208662 | *LEPR* | 1:65987164 | C/G | 23563609 | 0.139 [0.037, 0.242] | 7.66x10<sup>-03</sup> |
| rs6235 | *PCSK1* | 5:95728898 | G/C | 18604207 | 0.090 [0.023, 0.158] | 8.82x10<sup>-03</sup> |
| rs9356744 | *CDKAL1* | 6:20685486 | T/C | 22344219 | 0.071 [0.005, 0.137] | 0.035 |
| rs7988412 | *MTIF3* | 13:28000282 | T/C | 25673413 | 0.090 [0.005, 0.175] | 0.037 |
| rs1780050 | *NEXN* | 1:78400540 | A/C | 25673413 | 0.063 [0.002, 0.124] | 0.042 |
| rs526134 | *USP37* | 2:219402371 | G/A | 25673413 | 0.066 [0.000, 0.132] | 0.049 |
| rs980828 | *NOS1AP* | 1:162306415 | G/T | 25133637 | 0.050 [-0.010, 0.110] | 0.100 |
| rs17001561 | *SCARB2* | 4:77096118 | A/G | 25673413 | 0.070 [-0.017, 0.157] | 0.113 |

| **SNP** | **Gene** | **Chr:Position** | **E/O** | **PMID** | **$\beta_{OLS}$ [95%CI]** | **p-value** |
| --- | --- | --- | --- | --- | --- | --- |
| rs6232 | *PCSK1* | 5:95751785 | C/T | 18604207 | 0.095 [-0.041, 0.232] | 0.172 |
| rs749767 | *KAT8* | 16:31124407 | A/G | 25673413 | 0.042 [-0.022, 0.105] | 0.199 |
| rs1211166 | *NTRK2* | 9:87285992 | A/G | 23001569 | 0.041 [-0.034, 0.116] | 0.289 |
| rs2535633 | *ITIH4* | 3:52859630 | G/C | 24861553 | 0.024 [-0.037, 0.085] | 0.437 |
| rs10144353 | *PRKCH* | 14:61911157 | T/C | 23563609 | 0.044 [-0.067, 0.155] | 0.441 |
| rs1561288 | *ADCY3* | 2:25369002 | C/T | 23669352 | 0.024 [-0.047, 0.095] | 0.507 |
| rs2283228 | *KCNQ1* | 11:2849530 | C/A | 24861553 | -0.037 [-0.159, 0.085] | 0.550 |

Single or mixed interactions in the effects of SNPs that are not adjusted in regression models will produce variability in CQR estimates along the distribution of the outcome (see Supplemental Note). This variability can be detected and quantified using MR.^25,26^ Simulations showed that the power to detect such interactions using CQR and MR was not affected by the MAF or the main effects of the SNPs, but increased with the number of interactions as well as the main effects of the interacting covariate (see Supplemental Note and Figure S1). Yaghootkar, et al., have recently shown that differences in the prevalence of diseases outcomes (e.g. T2D) between sample and general populations can bias regression estimates of the main effects of SNPs on risk factors (e.g. BMI).^86^ However, the variability of CQR estimates across the sample distribution is not affected by biased main effects when CQR models are fitted with adjustment for disease status (see Supplemental Note). This was supported by simulations which showed that the prevalence of disease outcomes in sample populations had negligible effects on the power and Type-I error rate for detecting unadjusted interactions when CQR models were adjusted for disease status (see Supplemental Note and Figure S2).

MR models were fitted to assess the variability in the CQR estimates of BMI-associated SNPs along the sample distribution of BMI (Table 2, Figures 2 and S3). Significant positive associations (p<1.32×10^−03^) between BMI percentile and CQR estimates were detected for 9 of 37 SNPs (24%) including, *FTO* (rs1421085, *β_MR_* [95%CI]=0.49 [0.37, 0.62], p=8.69×10^−15^), *PCSK1* (rs6235, 0.32 [0.18, 0.46], 7.11×10^−06^), *TCF7L2* (rs7903146, 0.30 [0.17, 0.44], 9.60×10^−6^), *MC4R* (rs11873305, 0.60 [0.31, 0.89], 5.08×10^−05^), *FANCL* (rs12617233, 0.26 [0.13, 0.39], 5.30×10^−05^), *GIPR* (rs11672660, 0.29 [0.14, 0.45], 1.64×10^−04^), *MAP2K5* (rs997295, 0.23 [0.10, 0.35], 3.25×10^−04^), *FTO* (rs6499653, 0.25 [0.11, 0.40], 6.23×10^−04^) and *NT5C2* (rs3824755, 0.36 [0.15, 0.57], 7.90×10^−04^). The estimates from MR (*β_MR_*) quantify changes in the impact of each SNP on BMI across the sample distribution. For these 37 SNPs, the median *β_MR_* value [Q1, Q3] was 0.135 [0.094, 0.217] kg/m^2^ per Effect Allele per BMI Percentile. In this statistical framework *β_MR_* is equal to zero if all SNP interaction effects are also zero (see Supplemental Note). Positive *β_MR_* estimates indicate that effects of SNPs vary systemically by BMI percentile because unadjusted interactions are inflating the effects of SNPs in participants with high BMI.

**Table 2:** *Quantifying the effect of BMI percentile on conditional quantile regression (CQR) estimates using meta-regression (MR*). MR was used to model variability in the CQR estimates across BMI percentiles. Note that the percentiles were re-centered around the 50^th^ percentile so that the intercept from MR models corresponds to the main effect of the SNP at the median. (*) Denotes statistical significance at the Bonferroni adjusted threshold of p<1.32×10^−03^, RI_50_ is the re-centered intercept of the MR models, *β_MR_* is the effect of BMI percentile on CQR estimates (kg/m^2^ per Effect Allele per BMI Percentile), 95%CI are the 95% confidence intervals.

| SNP | Gene | $RI_{50}$ | $\beta_{MR}$ [95%CI] | p-value |
| --- | --- | --- | --- | --- |
| rs1421085 | *FTO* | 0.473 | 0.495 [0.370, 0.620] | $8.69 \times 10^{-15}$ * |
| rs6235 | *PCSK1* | 0.078 | 0.320 [0.180, 0.459] | $7.11 \times 10^{-06}$ * |
| rs7903146 | *TCF7L2* | 0.144 | 0.303 [0.169, 0.437] | $9.60 \times 10^{-06}$ * |
| rs11873305 | *MC4R* | 0.344 | 0.603 [0.311, 0.895] | $5.08 \times 10^{-05}$ * |
| rs12617233 | *FANCL* | 0.129 | 0.261 [0.134, 0.387] | $5.30 \times 10^{-05}$ * |
| rs11672660 | *GIPR* | 0.227 | 0.294 [0.141, 0.447] | $1.64 \times 10^{-04}$ * |
| rs997295 | *MAP2K5* | 0.131 | 0.228 [0.103, 0.352] | $3.25 \times 10^{-04}$ * |
| rs6499653 | *FTO* | 0.121 | 0.253 [0.108, 0.398] | $6.23 \times 10^{-04}$ * |
| rs3824755 | *NT5C2* | 0.222 | 0.362 [0.151, 0.574] | $7.90 \times 10^{-04}$ * |
| rs7553158 | *TNNI3K* | 0.099 | 0.196 [0.071, 0.322] | $2.12 \times 10^{-03}$ |
| rs10767664 | *BDNF* | 0.247 | 0.217 [0.064, 0.370] | $5.50 \times 10^{-03}$ |
| rs4788099 | *SH2B1* | 0.151 | 0.194 [0.057, 0.332] | $5.59 \times 10^{-03}$ |
| rs17066846 | *MC4R* | 0.124 | 0.215 [0.063, 0.367] | $5.61 \times 10^{-03}$ |
| rs9356744 | *CDKAL1* | 0.063 | 0.186 [0.050, 0.322] | $7.35 \times 10^{-03}$ |
| rs6453133 | *HMGCR* | 0.130 | 0.177 [0.040, 0.314] | 0.011 |
| rs2819347 | *LMOD1* | 0.111 | 0.137 [0.004, 0.269] | 0.044 |
| rs2075650 | *TOMM40* | 0.283 | 0.161 [-0.019, 0.341] | 0.079 |
| rs4946932 | *FOXO3* | 0.106 | 0.120 [-0.016, 0.256] | 0.084 |
| rs2984618 | *TAL1* | 0.069 | 0.108 [-0.019, 0.235] | 0.095 |
| rs980828 | *NOS1AP* | 0.024 | 0.095 [-0.030, 0.220] | 0.135 |
| rs1788826 | *NPC1* | 0.109 | 0.094 [-0.036, 0.224] | 0.156 |
| rs11570094 | *SPI1* | 0.103 | 0.096 [-0.039, 0.231] | 0.163 |
| rs7988412 | *MTIF3* | 0.088 | 0.109 [-0.062, 0.280] | 0.212 |
| rs2283228 | *KCNQ1* | 0.003 | 0.147 [-0.094, 0.388] | 0.232 |
| rs739564 | *IQCK* | 0.122 | 0.100 [-0.065, 0.265] | 0.234 |
| rs526134 | *USP37* | 0.062 | 0.079 [-0.055, 0.212] | 0.247 |
| rs2272903 | *TFAP2B* | 0.145 | 0.113 [-0.084, 0.310] | 0.261 |
| rs2836754 | *ETS2* | 0.086 | 0.073 [-0.060, 0.206] | 0.280 |
| rs2535633 | *ITIH4* | 0.016 | 0.068 [-0.059, 0.194] | 0.296 |
| rs11208662 | *LEPR* | 0.142 | 0.111 [-0.105, 0.327] | 0.314 |

| **SNP** | **Gene** | **RI<sub>50</sub>** | **$\beta_{MR}$ [95%CI]** | **p-value** |
| --- | --- | --- | --- | --- |
| rs6232 | *PCSK1* | 0.075 | 0.133 [-0.137, 0.404] | 0.334 |
| rs749767 | *KAT8* | 0.048 | 0.058 [-0.075, 0.191] | 0.390 |
| rs1561288 | *ADCY3* | 0.027 | -0.037 [-0.185, 0.112] | 0.627 |
| rs10144353 | *PRKCH* | 0.043 | 0.049 [-0.171, 0.269] | 0.662 |
| rs1211166 | *NTRK2* | 0.029 | -0.027 [-0.179, 0.126] | 0.731 |
| rs17001561 | *SCARB2* | 0.068 | -0.020 [-0.194, 0.154] | 0.824 |
| rs1780050 | *NEXN* | 0.045 | 0.010 [-0.117, 0.136] | 0.883 |

Since height is known to be highly heritable, analyses were extended to height as a reference to the BMI results.^22,87,88^ OLS models were fitted for each of 125 height-associated SNPs and all but two were verified to increase height (Table S6). CQR and MR were used to estimate variation in the effects of these SNPs on height as above (Figure S4 and Table S7). Only one height-associated SNP, *IGF1* (rs6219, *β_MR_* [95%CI]=0.48 [0.23, 0.73], p=1.80×10^−4^), showed significantly (p<3.97×10^−4^) increased effects along the sample height distribution (Table S8). For height-associated SNPs, the median *β_MR_* value [Q1, Q3] was 0.002 [−0.056, 0.085] cm per Effect Allele per Height Percentile. Thus, CQR estimates for height-associated SNPs were predominantly consistent across height percentiles and <1% showed evidence of unadjusted interactions, compared to 24% of BMI/obesity-associated SNPs.

BMI/obesity- and height-associated SNPs were combined into gene scores (GS-BMI and GS-Height, respectively) to examine the overall association of these SNPs across the sample distribution. OLS models were used to verify the positive association between GS-BMI and GS-Height with their respective traits (Table 3). CQR models for GS-BMI showed steadily increasing effects with increasing percentiles, while CQR models for GS-Height did not vary across percentiles (Figure 3). MR analysis indicated that percentiles were significantly and positively associated with CQR estimates for GS-BMI (*β_MR_* [95%CI]=0.15 [0.13, 0.17], 7.03×10^−37^) but not GS-Height (0.01 [−0.01, 0.02], 0.499) (Table 3). At the 10^th^ and 90^th^ BMI percentiles, each additional effect allele of GS-BMI increased BMI by 0.054 and 0.167 kg/m^2^ (3.1-fold increase), respectively; while each additional allele of GS-Height increased height by 0.172 and 0.180, respectively (Table S5 and S7). Thus, in 1.73m tall persons at the 10^th^ BMI percentile, carrying 10 additional BMI-increasing alleles was associated with 1.6 kg of extra weight, while at the 90^th^ BMI percentile this was associated with 5.0 kg of extra weight. Furthermore, at the 10^th^ and 90^th^ BMI percentiles, the proportion of trait variance explained by GS-BMI increased (2.7 fold, 0.130% to 0.357%), while that of GS-Height was stable (1.825% to 1.822%) (Tables S5 and S7). These results support the conclusion that the impact of BMI-associated SNPs was larger for individuals with high BMI, which contrasts with the impact of height-associated SNPs which varied little by height.

**Figure 3:**
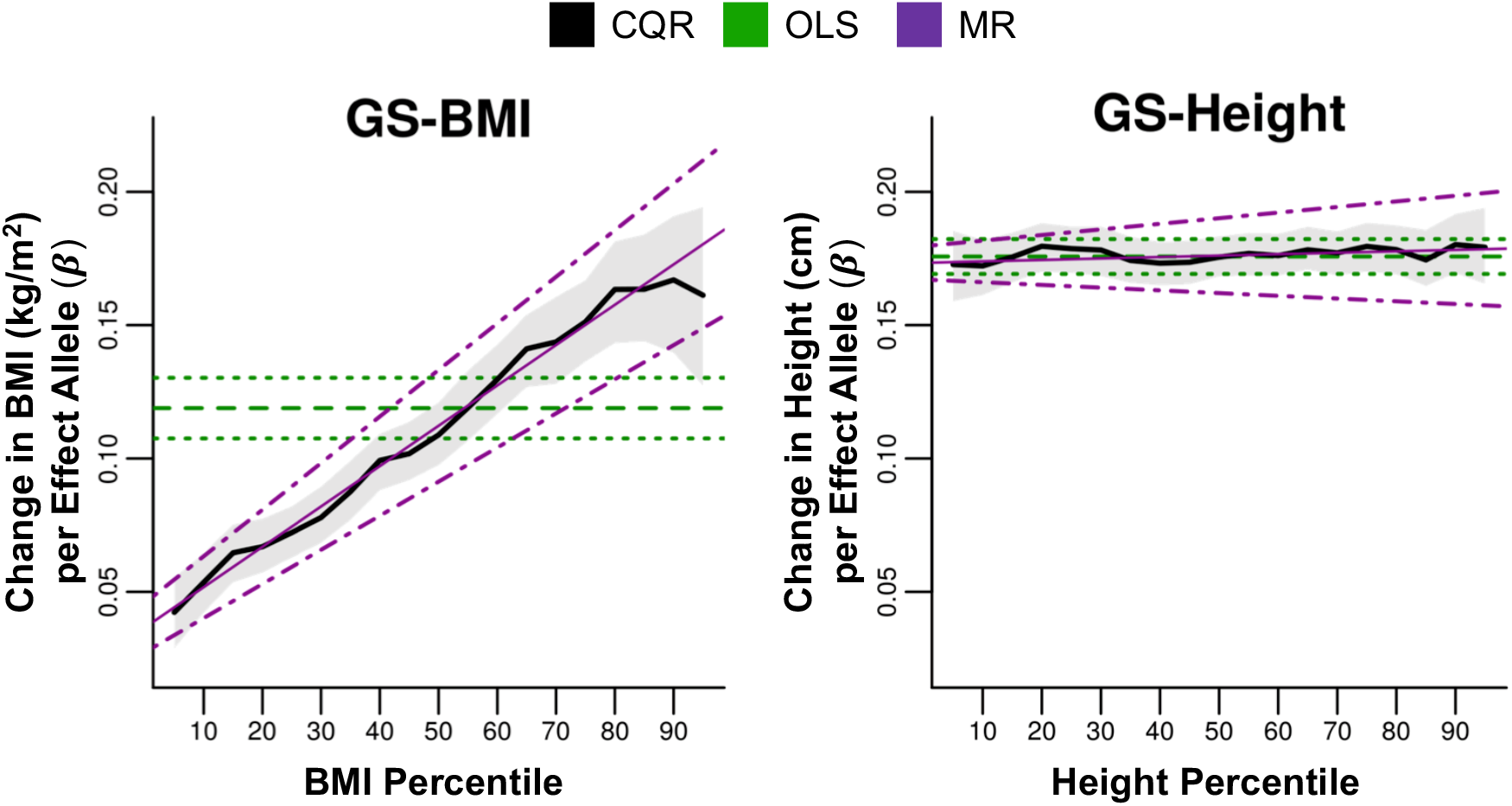
*The effects of GS-BMI and GS-Height across the sample distribution of BMI and height, respectively*. As in Figure 2, CQR models of the GS-BMI and GS-Height were plotted against the BMI percentile and height percentile, respectively. The thick-black line is the estimated change in each trait per effect allele (GS-BMI, *β_CQR_*, kg/m^2^ per Effect Allele; GS-Height, *β_CQR_*, cm per Effect Allele) and shaded-grey region represents the 95% confidence intervals. Also plotted are the OLS regression estimates (GS-BMI, *β_OLS_* in kg/m^2^ per Effect Allele; GS-Height, *β_OLS_*, cm per Effect Allele, horizontal-dashed-green line) and 95% confidence intervals (horizontal-dotted-green lines). The change in CQR estimates across outcome percentiles was modeled using meta-regression (MR). Estimates from MR (GS-BMI, *β_MR_*, kg/m^2^ per Effect Allele per BMI Percentile; GS-Height, *β_MR_*, cm per Effect Allele per Height Percentile; thin-magenta line) and the 95% confidence intervals (dotdashed-magenta lines) were also plotted.

**Table 3:**
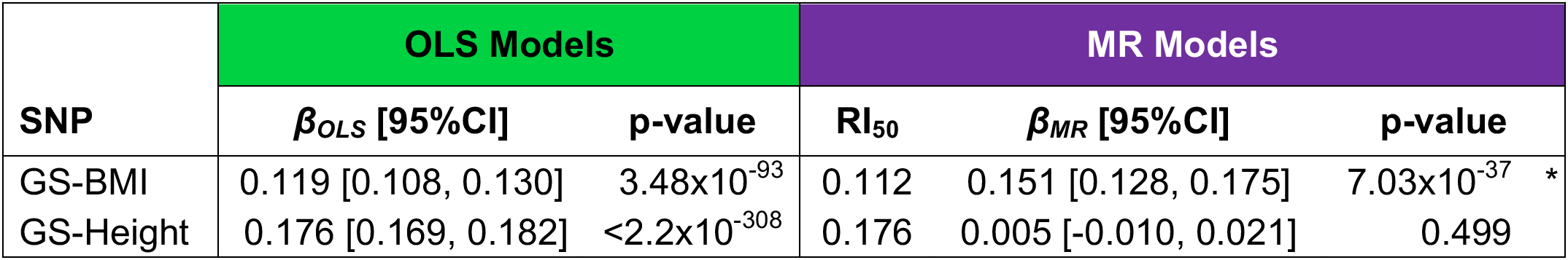
*Analysis of the GS-BMI and GS-Height*. BMI/Obesity- and height-associated SNPs were combined into gene scores (GS-BMI and GS-Height, respectively). As in Table 1, the results from ordinary least squares (OLS) models are presented. Furthermore, as in Table 2, meta-regression (MR) analysis was applied to quantify the effects of trait (BMI and height) percentile on the conditional quantile regression (CQR) estimates for GS-BMI and GS-Height, respectively. (*) Denotes statistical significance at the Bonferroni adjusted threshold of p<1.32×10^−03^ for GS-BMI and p<3.97×10^−04^ for GS-Height. *β_OLS_* is the effect size (GS-BMI, kg/m^2^ per Effect Allele; GS-Height, cm per Effect Allele) from OLS Models, RI_50_ is the re-centered intercept of the MR models (same units as *β_OLS_*), *β_MR_* is the effect size (GS-BMI, kg/m^2^ per Effect Allele per BMI Percentile; GS-Height, cm per Effect Allele per Height Percentile) from MR models, and 95%CI are the 95% confidence intervals.

Excluding 7 SNPs that have also been associated with comorbidities of obesity from the GS, GS-BMI (Stringent), did not alter the pattern of increasing effects across the sample BMI distribution (Figure S5).^48–55^ Moreover, MR analysis indicated that BMI percentile was significantly and positively associated with the CQR estimates for GS-BMI (Stringent) (*β_MR_* [95%CI]=0.14 [0.11, 0.16], p=2.18×10^−23^). In addition, CQR models were refitted with adjustment for diabetic status as this was shown to mitigate the effects of possible stratification within the sample population (see Supplemental Note and Figure S2). Of the 9 SNPs whose effects showed significant increases across the sample BMI distribution (Table 2 and Figure 2), 3 have also been associated with glucose homeostasis and T2D, namely *GIPR* (rs11672660), *TCF7L2* (rs7903146) and *PCSK1* (rs6235).^48,50–53^ Refitting CQR models with adjustment for diabetic status had little impact on the results from MR analysis of these SNPs or GS-BMI (Table S9). To address the potential effects of scaling, analyses were conducted on 1) transformed BMI and 2) the residuals after adjusting for the median effects of each SNP. The scaling effect refers to potential correlations between mean and variance effects that result from skewness.^63^ Transformations counteract scaling by reducing skewness, which may suppress some potentially useful distributional information, while residual analysis addresses scaling directly to preserve distributional information (Figure S6). Despite a reduction in sensitivity, significant associations between log-transformed BMI percentile and CQR estimates were detected for 4 of 9 SNPs including *FTO* (rs1421085), *PCSK1* (rs6235), *TCF7L2* (rs7903146), *FANCL* (rs12617233) and GS-BMI (Table S9). Rank transformation of BMI further reduced the overall sensitivity and only *PCSK1* (rs6235) and GS-BMI continued to show significant associations, while most of the remaining SNPs were relegated to nominal levels of significance (Table S9). Analysis of SNP-adjusted BMI residuals detected significant associations between BMI percentile and CQR estimates for all 9 SNPs and GS-BMI (Table S9). Additional sensitivity analysis that included modelling the effects of age linearly or testing fewer percentiles (i.e. every 10^th^ percentile from the 5^th^ to 95^th^ BMI percentiles) also showed no substantial changes to MR results (Table S9). Furthermore, calculating the GS for each trait without imputing missing genotypes did not affect results for GS-BMI and GS-Height (Figure S5). Finally, the results from CQR were compared to those from conventional subgroup analysis. To this end, the effects of genotype on the risk of OW, Ob-l, Ob-ll and Ob-Ill was evaluated separately using logistic regression (Table S10). The odds ratios (ORs) of each SNP for each category were plotted against the BMI percentiles of the corresponding category and CQR estimates were then overlaid on these bar plots. The patterns from logistic regression models across BMI categories were qualitatively consistent with the patterns from CQR models at comparable BMI percentiles (Figure S7).

## DISCUSSION

The aim of this study was to investigate variations in the impact of 37 BMI/obesity-associated SNPs across the distribution of BMI. We introduced a method that applies CQR to model the effects of SNPs at different percentiles of the sample BMI distribution and then estimated variability in these effects using MR. CQR estimates at different percentiles were shown to be uniform if all unadjusted SNP interactions are zero (see Supplemental Note). It follows that SNPs whose CQR estimates vary significantly across the sample BMI distribution are regulated by such interactions.

CQR analysis revealed distinct profiles of associations of BMI/obesity SNPs across the sample BMI distribution. Several of these SNPs had effects that increased steadily at higher BMI percentiles while others had uniform effects that varied little across BMI percentiles (Figures 2 and S3). One other study has used CQR to explore the relationship between genetic variants and BMI in a modest sample of adults for *FTO* (rs1558902) and a GS.^89^ The patterns reported by that study are consistent with the results reported here.^89^ Two other reports used CQR to investigate the effects of SNPs on BMI in European children and their results are also comparable with those here.^90,91^ Overall, the high degree of correspondence between previously reported CQR results with European children and those from adults presented here emphasizes the robustness of these findings. Furthermore, the patterns observed using CQR analysis were compared to those from conventional logistic regression (subgroup analysis) as Berndt et al., has demonstrated that the genetic architecture of BMI overlaps strongly with BMI categories (Table S10).^92^ The patterns across BMI categories from logistic regression were largely consistent with those from CQR (Figure S7). CQR overcomes several of the limitations of subgroup analysis as it utilizes the entire sample data to estimate regression parameters on the same scale as the continuous outcome and comparing CQR estimates from different quantiles is relatively intuitive and easy.^23,92^

MR was applied to model changes in the effects of BMI/obesity SNPs across the sample BMI distribution.^25,26^ Results from MR showed that BMI percentile was positively and significantly associated with CQR estimates for 9 of 37 SNPs (24%). In addition, nominal associations were also observed for several other SNPs and the median *β_MR_* [Q1, Q3] was 0.135 [0.094, 0.217] kg/m^2^ per Effect Allele per BMI Percentile (Table 2 and Figure S3). This is supported by the analysis of GS-BMI which also showed significantly increasing effects across the sample BMI distribution (Figure 3 and Table 3). These findings indicate that unadjusted interactions enhanced the effects of BMI-associated SNPs at higher BMI levels. Modelling the effects of age linearly or considering fewer BMI percentiles (every 10^th^ rather than every 5^th^ percentile) had minimal effects on these results (Table S9). Although CQR does not make any assumptions about the outcome distribution, BMI is often transformed (log and inverse-rank) to accommodate the normality assumption of OLS at the cost of suppressing some distributional information. Transformations disproportionately compress distances between samples to impose symmetry on distributions and some have argued that rank transformation in particular is overly conservative.^63,93,94^ Part of the novelty of our approach is that CQR and MR leverage precisely this distributional information to extract evidence of gene interactions and we expect transformations that suppress distances between ranked samples to reduce the sensitivity of this method. Despite decreased sensitivity, significant associations between BMI percentile and CQR estimates were detectable with log transformed BMI for 4 of the 9 SNPs and GS-BMI while the rest showed nominal associations (Table S9). Rank-transformation decreased sensitivity even further and only *PCSK1* (rs6235) and GS-BMI had significant detectable associations, while many of the remaining 9 SNPs showed nominal associations.

Scaling refers to the phenomenon where the mean and variance effects of a SNP can be correlated when the outcome distribution is skewed.^63^ This is typically addressed using transformations to impose symmetry on the outcome distribution. Although quantile based methods do not rely on the mean and variance they may also be susceptible to scaling effects (Figure S6).^24^ To examine the possible role of scaling in CQR and MR without compromising sensitivity, we conducted analyses on the residuals after adjusting for the median effects of each SNP. Residual analysis was shown to effectively mitigate scaling effects by reducing the correlation between SNP main effects and *β_MR_* (Figure S6). Significant associations between residual BMI percentile and CQR estimates were detected for all 9 SNPs and GS-BMI, indicating that the associations persisted even after adjusting for SNP main effects (Table S9). These results confirm that scaling effects did not substantially contribute to our findings. Future work aimed at better understanding the phenomenon of scaling would be useful for analysis of quantitative traits with asymmetric distributions.

There is evidence that differences in disease prevalence (e.g. T2D) between sample and general populations can result in the stratification of secondary traits (e.g. BMI) that are risk factors for disease.^86^ This stratification can compromise regression estimates of the main effects of SNPs on secondary traits and naively adjusting regression models for disease status may not adequately address this.^86^ While the main effects of SNPs from disease-adjusted regression models are susceptible to stratification bias, the variation of SNP effects across the sample distribution is not (see Supplemental Note). This is evident in simulations which showed that stratification had little effect on the power and Type-1 error rate of MR analysis when CQR models were adjusted for disease status (Figure S2). As *GIPR* (rs11672660), *TCF7L2* (rs7903146) and *PCSK1* (rs6235) have been associated with glucose homeostasis and T2D, CQR models were refitted with adjustment for diabetic status and analyzed using MR.48,5o,53 jhese SNPs and the GS continued to show significantly increasing effects across the sample BMI distribution with this adjustment, which demonstrated that the results were not an artifact of possible sample stratification (Table S9). Although estimating the variability of disease-adjusted CQR estimates across the sample distribution using MR is robust against stratification bias, future studies aimed at estimating the main effects of SNPs using CQR should implement methods to address this potential source of bias.^95^ A total 7 of the 37 obesity predisposing loci that were selected for analysis have also been associated with comorbidities of obesity including glucose homeostasis, T2D, lipid levels and CRP levels.^48–55^ Excluding these SNPs from the GS did not alter the pattern observed across the sample BMI distribution or affect the results from MR analysis, which suggested that these findings do not stem from the influence of comorbidities at high BMI levels (Figure S5).

Although BMI was the primary focus of this report these analyses were also applied to height. This was important because analysis of height could shed light on the nature of the unadjusted interactions that were detected. BMI is a composite of both height and weight, where height is one of the most heritable complex human traits and weight is strongly influenced by environmental exposures and behavior.^11,96^ If unadjusted interactions in the effects of BMI/obesity-associated SNPs are predominantly due to GXG interactions, then it is reasonable to suppose that they would be detected with a similar frequency in other quantitative traits such as height. On the other hand, if GXE interactions predominate then they may be less frequently detected in quantitative traits with a smaller environmental component (i.e. height). CQR models for 125 height-associated SNPs were mostly uniform and exhibited little variability across height percentiles (Figure S4). Only one significant association between height percentiles and CQR estimates for height SNPs was detected by MR and the median *β_MR_* [Q1, Q3] was 0.002 [−0.056, 0.085] cm per Effect Allele per Height Percentile (Table S8). Moreover, the effects of GS-Height did not vary along the sample height distribution, which suggests that unadjusted interactions do not impact the genetic architecture of height to same extent that they do for BMI (Table 3 and Figure 3). The simplest explanation for the discrepancy between the results for GS-BMI and GS-Height is that the unadjusted interactions detected from GS-BMI are predominantly GXE interactions. It is important to consider that gene interactions described here include variants with non-linear genetic models of effects where the presence of one allele changes the effects of a second allele.^57–58^

GXE interactions for SNPs in the *FTO* gene have been reported for physical activity, food intake, dietary salt, alcohol consumption and sleep duration.^97–100^ In addition, the effects of *TCF7L2* (rs12255372) on BMI showed interactions with fat intake in a weight-loss trial.^101^ Our analyses also pointed to significant interactions for *FTO* (rs1421085) and *TCF7L2* (rs7903146), but suggested that such interactions may extend to additional BMI/obesity-associated SNPs including *PCSK1* (rs6235), *MC4R* (rs11873305), *FANCL* (rs12617233), *GIPR* (rs11672660), *MAP2K5* (rs997295), *FTO* (rs6499653), *NT5C2* (rs3824755) and GS-BMI. This is entirely consistent with a report showing that the effects of GS-BMI (29 SNPs) was increased by greater exposure to obesogenic environments and another demonstrating interactions between GS-BMI (69 SNPs) and several obesogenic drivers including socioeconomic status, TV watching, ‘westernized’ diets and physical activity.^14,102^ These reports also support the argument that the unadjusted interactions detected for BMI SNPs are predominately GXE interactions. Environmental modification of the effects of genetic variants raises the possibility that preventive measures, sustained lifestyle modifications and therapeutic interventions may attenuate some of the genetic elements of BMI. Indeed, the overall effect of BMI/obesity SNPs is minimal at low BMI levels (Figures 2 and 3). If weight-gain leads to a genetically driven ‘vicious circle’, then weight-loss can lead to a genetically driven ‘virtuous circle’. Investigating additional BMI-associated SNPs using CQR and MR to uncover the full extent of unadjusted interactions in the architecture of BMI will be the focus of future studies.

This study is the largest yet to apply CQR to examine how the effects of SNPs vary with BMI and establishes quantitative support for hitherto qualitative descriptions of CQR. The combined utility of CQR and MR presents a contemporary statistical framework to cue hypotheses on gene interactions, better define clinical risks associated with genetic profiles and prioritize clinical targets. Future studies aimed at distinguishing variants whose effects are modified by unadjusted interactions from those with fixed effects could advance the field of precision medicine. With the combined application of CQR and MR, this can now be achieved solely using information contained within the sample outcome distribution.

## SUPPLEMENTAL DATA

Supplemental data include one Supplemental Note, six figures and ten tables

## ACKNOWLEDGEMENTS

We thank Aihua Li for her assistance in database management and Guillaume Pare for his critical review of the manuscript. Sebastien Robiou-du-Pont was supported by the Heart and Stroke Foundation of Ontario. Sonia S. Anand holds the Heart and Stroke Foundation of Ontario Michael G. DeGroote endowed Chair in Population Health and a Canada Research Chair in Ethnicity and Cardiovascular Disease, Hertzel C. Gerstein holds the Aventis PHRI Chair in Diabetes, Salim Yusuf holds the Heart and Stroke Endowed Chair in Cardiovascular Research, and David Meyre holds a Canada Research Chair in Genetics of Obesity. Further acknowledgements are presented in the Supplemental Note.

## WEB RESOURCES

The URLs for databases and software resources described here are:

dbGaP: https://www.ncbi.nlm.nih.gov/gap

1000g: http://www.internationalgenome.org/

R statistical software: http://www.r-project.org/

PLINK: https://www.cog-genomics.org/plink2/

GWAS Catalog: https://www.ebi.ac.uk/gwas/

